# *P. falciparum* gametocyte density and infectivity in peripheral blood and skin tissue of naturally infected parasite carriers

**DOI:** 10.1101/803296

**Authors:** Elamaran Meibalan, Aissata Barry, Matthew P. Gibbins, Shehu Awandu, Lisette Meerstein-Kessel, Fiona Achcar, Selina Bopp, Christopher Moxon, Amidou Diarra, Siaka Debe, Nicolas Ouédraogo, Ines Barry-Some, Emilie Badoum, Traoré Fagnima, Kjerstin Lanke, Bronner P. Gonçalves, John Bradley, Dyann Wirth, Chris Drakeley, Wamdaogo Moussa Guelbeogo, Alfred B. Tiono, Matthias Marti, Teun Bousema

## Abstract

Transmission of *Plasmodium falciparum* depends on the presence of mature gametocytes that can be ingested by mosquitoes taking a bloodmeal when feeding on human skin. It has long been hypothesised that skin sequestration contributes to efficient transmission. Although skin sequestration would have major implications for our understanding of transmission biology and the suitability of mosquito feeding methodologies to measure the human infectious reservoir, it has never been formally tested. In two populations of naturally infected gametocyte carriers from Burkina Faso, we assessed transmission potential to mosquitoes and directly quantified male and female gametocytes and asexual parasites in: i) finger prick blood, ii) venous blood, iii) skin biopsies, and in pools of mosquitoes that fed iv) on venous blood or, v) directly on the skin. Whilst more mosquitoes became infected when feeding directly on the skin compared to venous blood, concentrations of gametocytes in the subdermal skin vasculature were identical to that in other blood compartments. Asexual parasite densities, gametocyte densities and sex ratios were identical in the mosquito blood meals taken directly from the skin of parasite carriers and their venous blood.

We also observed sparse gametocytes in skin biopsies from legs and arms of gametocyte carriers by microscopy. Taken together, we provide conclusive evidence for the absence of significant skin sequestration of *P. falciparum* gametocytes. Gametocyte densities in peripheral blood are thus informative for predicting onward transmission potential to mosquitoes. Quantifying this human malaria transmission potential is of pivotal importance for the deployment and monitoring of malaria elimination initiatives.

**IMPORTANCE:** Our observations settle a long-standing question in the malaria field and close a major knowledge gap in the parasite cycle. By deploying mosquito feeding experiments and stage-specific molecular and immunofluorescence parasite detection methodologies in two populations of naturally infected parasite carriers, we conclusively reject the hypothesis of gametocyte skin sequestration. Our findings provide novel insights in parasite stage composition in human blood compartments, mosquito bloodmeals and their implications for transmission potential. We demonstrate that gametocyte levels in venous or finger prick blood can be used to predict onward transmission potential to mosquitoes. Our findings thus pave the way for methodologies to quantify the human infectious reservoir based on conventional blood sampling approaches to support the deployment and monitoring of malaria elimination efforts for maximum public health impact.

## INTRODUCTION

Significant reductions in malaria burden in recent decades have stimulated malaria elimination initiatives (1). It is widely accepted that malaria elimination with current tools is unlikely for the majority of African settings (2). Therefore, novel interventions are needed and approaches that specifically reduce malaria transmission may be of key importance (3). Transmission of malaria depends on the presence of mature male and female gametocytes that circulate in the bloodstream and may be ingested by mosquitoes from the subdermal capillaries upon blood feeding. For *P. falciparum*, these circulating mature gametocytes are the product of a prolonged developmental process that starts with commitment of asexual parasites to the sexual pathway upon activation of AP2-G(4, 5). Developing gametocytes are sequestered for 10-12 days, primarily to the bone marrow and spleen (6), until their release into the blood circulation as mature gametocytes. Mosquitoes may become infected when feeding and ingesting mature male and female gametocytes, even if their densities in the peripheral blood are low (7). Interestingly, mosquito infections have been observed from gametocyte donors whose low gametocyte density appears incompatible with transmission (8). Mosquito infection rates are typically higher when mosquitoes feed directly on the skin of gametocyte carriers, as compared to feeding on venous blood through an artificial membrane (9, 10). In addition to a strategic adjustment of gametocyte sex-ratio to maximize transmission success (7, 11, 12), gametocyte aggregation and sequestration may facilitate mosquito infections from low gametocyte densities. Aggregation of gametocytes in blood meals has been observed (13) and may increase the chance that both male and female gametocytes are ingested. Gametocyte sequestration in the skin tissue may further increase transmission rates and would parallel sequestration patterns for other human parasites. The importance of skin sequestration for transmission to invertebrate vectors was recently demonstrated for skin-dwelling *Trypanosoma brucei* (14), as was previously reported for *Onchocerca volvulus,* different species of *Mansonella*, *Leishmania infantum* and *L. donovani*, where parasite burden in the skin is the best predictor of infectiousness (15–18).

Indirect evidence for skin sequestration of mature gametocytes in the microvasculature of the skin was first described following surveys conducted in the 1940s and 1950s in DR Congo: gametocyte prevalence in a survey using skin scarification was 3-fold higher compared to a survey 5 years earlier using finger prick blood (19). In a follow up study with 1243 paired samples, a more modest 13.4% increase in *P. falciparum* parasite prevalence and 15.6% increase in gametocyte prevalence was observed when blood and dermal fluids from skin scarification were used for sample preparation instead of finger prick blood (20). The hypothesized skin sequestration of intra-erythrocytic *P. falciparum* gametocytes may be related to mechanical retention in cutaneous capillaries (21, 22), analogous to the reversible rigidity that likely prevents immature gametocytes from entering circulation (23, 24). Alternatively, sequestration may be related to gametocyte cytoadhesive properties (25) mediated by parasite proteins that are present on the infected red blood cell (iRBC) surface, analogous to adhesion of asexual *P. falciparum* parasites to receptors on human vascular endothelial cells by *P. falciparum* erythrocyte membrane-1 (PfEMP1)(26).

Whilst sequestration of mature gametocytes in the skin of naturally infected individuals remains speculative, it may play an important role in determining *Plasmodium* transmission efficiency (8, 22). Here, we report on two independent studies in naturally infected gametocyte carriers from Burkina Faso where we quantified mature *P. falciparum* gametocytes in skin tissue, blood samples and mosquito blood meals in association with onward transmission to *Anopheles* mosquitoes.

## RESULTS

A total of 31 individuals aged 15-48 (median 29) participated in experiments with paired skin feeding (27) and membrane feeding (28). The median number of dissected mosquitoes per experiment was 35 (interquartile range (IQR) 33-37) for direct skin feeding and 73 (IQR 69-82) for membrane feeding. Of 31 paired experiments, 18 (58.1%) direct skin feeding and 22 (71.0%) membrane feeding experiments resulted in at least one infected mosquito (p=0.289). Total gametocyte density, quantified in venous blood by quantitative reverse transcriptase PCR (qRT-PCR) targeting female-specific *Pfs25* mRNA and male-specific *Pfmget* mRNA (29), was positively associated with the proportion of mosquitoes that became infected following direct skin feeding (Spearman ρ=0.415, p=0.0204) or membrane feeding (Spearman ρ=0.596, p = 0.0004) (Figure 1A). The proportion of infected mosquitoes was higher by direct skin feeding as compared to membrane feeding assays (odds ratio 2.01; 95% CI 1.21 – 3.33, p = 0.007), in line with previous studies (9, 10, 30). The medium number of oocysts was 4 (IQR 2-7.5; maximum 38) for mosquitoes that became infected after feeding directly on the skin and 2 (IQR 1-5; maximum 24) for mosquitoes that became infected after feeding on venous blood through a membrane feeder.

**Figure 1.**
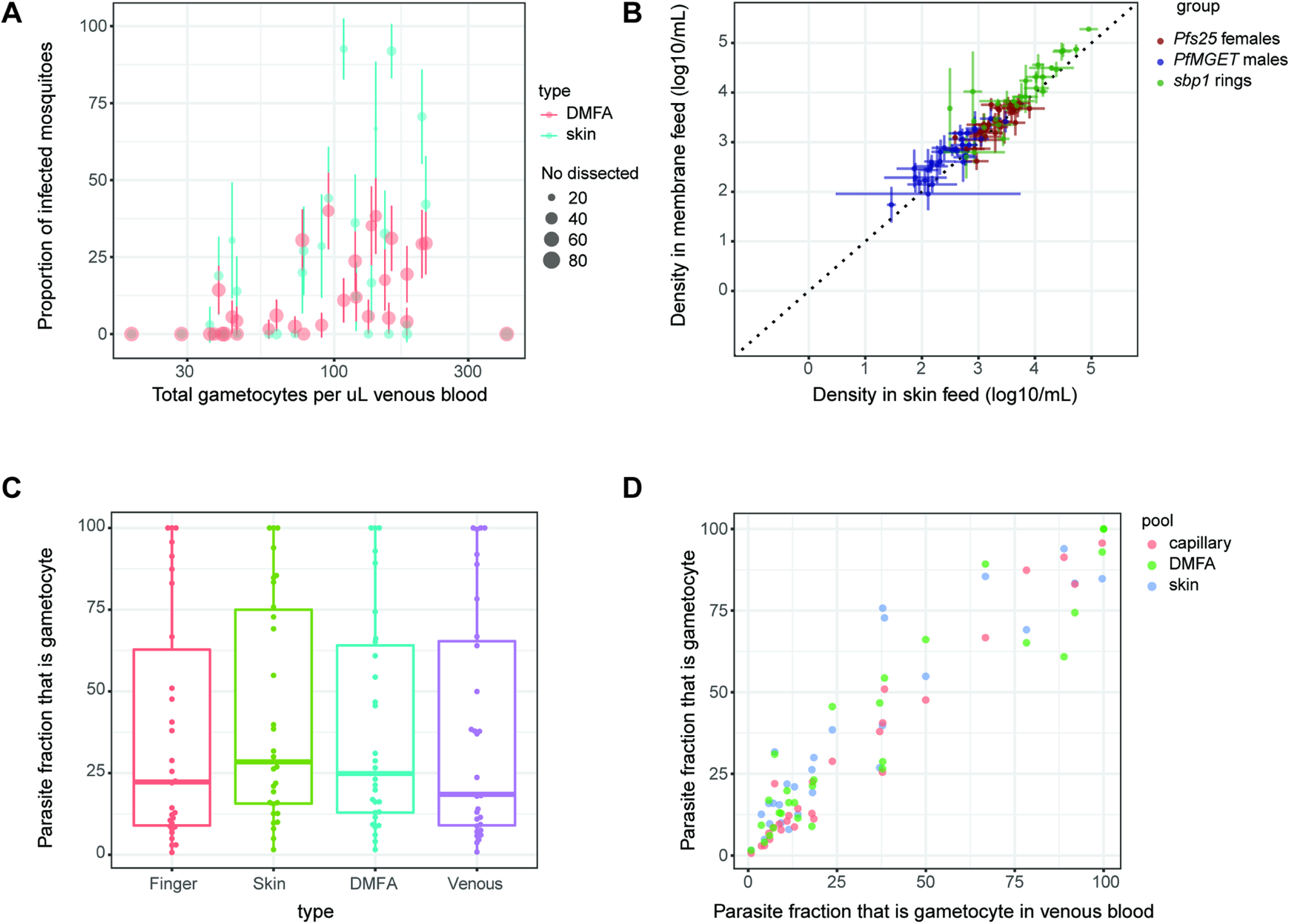
The density and infectivity of gametocytes in different blood compartments. **A.** Gametocyte density in venous blood in association with the proportion of mosquitoes that become infected when feeding directly on the skin of the blood donor (blue) or on venous blood offered through an artificial membrane feeder (red). Size indicates the number of examined mosquitoes; error bars indicate the 95% confidence interval around the proportion of infected mosquitoes. **B.** The density of ring stage asexual parasites (green), male gametocytes (blue) and female gametocytes (red) in mosquito blood meals when feeding directly on the skin (X-axis) versus venous blood offered through an artificial membrane feeder (Y-axis). Error bars indicate the standard error of density estimates in pools of mosquitoes fed directly on the skin (median 3 pools) or venous blood (median 4 pools). **C.** The fraction of the total parasite biomass that is gametocyte in finger prick capillary blood (red), mosquitoes that fed directly on the skin (green), mosquitoes that fed on venous blood (blue) or venous blood (purple). The box plot indicates median, interquartile range and range; dots indicate individual samples. **D.** The gametocyte fraction in venous blood (X-axis) versus on the Y-axis finger prick capillary blood (red; Spearman ρ=0.970; p<0.0001), mosquitoes that fed directly on the skin (green; Spearman ρ= 0.916; p<0.0001), mosquitoes that fed on venous blood (green; Spearman ρ=0.912; p<0.0001).

To examine whether this higher infectivity in direct skin feeding assays was related to higher ingested gametocyte densities, or to a higher gametocyte fraction in the blood meal, we directly quantified gametocytes and asexual parasites in mosquito blood meals. The blood content of individually fed mosquitoes was released into an RNA preservative 15 minutes after starting the feeding; RNA was then extracted and quantified from pools of 4 mosquitoes. We quantified asexual parasites by skeleton-binding protein 1 *sbp1* qRT-PCR (31) and gametocytes (*Pfs25* and *Pfmget* qRT-PCR) in a median of 3 mosquito pools per participant, each containing 4 individual mosquitoes, from skin-feeding (range=2-3) and 4 pools per participant, each containing 4 individual mosquitoes, from membrane feeding (range=2-4). We observed strong correlations between parasite quantities in pools of mosquitoes that fed on skin or venous blood through artificial membranes for asexual ring-stage parasites (r=0.921, p<0.0001), male (r=0.790, p<0.0001) and female gametocytes (r=0.655, p=0.0001) (Figure 1B). We also expressed gametocytes as a fraction of the total parasite biomass. This fraction ranged from very low (<1% gametocytes in an individual with 21,086 ring-stage asexual parasites/µL and 179 gametocytes/µL) to 100% in 3 individuals without asexual parasites detected by qRT-PCR (Figure 1C). We observed no tendency towards a higher fraction of gametocytes in skin-fed mosquitoes or capillary blood compared to venous blood (Figure 1D).

In a complementary study, 9 adult gametocyte carriers participated in skin biopsy sampling. After a screening visit, participants were seen on 2 occasions spaced 4 days apart. One participant came on day 5 for the return visit instead of day 4; one other participant withdrew consent prior to the second visit. On each occasion, venous blood, finger prick blood and 4 small skin biopsy punches were taken from the leg (n=2) and arm (n=2). Half of these biopsies were used for RNA extraction; the other half for histological assessments. Male and female gametocytes and ring-stage asexual parasites were quantified by qRT-PCR to calculate the gametocyte fraction in finger prick blood (16 observations; 9 donors), venous blood (n=16; 9 donors), as well as skin tissue from the arm (n=13; 7 donors) and leg (n=12; 8 donors). Gametocytes were detected in all tissue and all blood samples by qRT-PCR; asexual parasites were detected in 17/25 tissue and in 30/32 blood samples. The gametocyte fraction was highly variable between donors (and between time-points) whilst estimates from the different compartments from the same donor and time-point showed strong correlation: the gametocyte fraction in venous blood was strongly associated with that in finger prick blood (Spearman ρ =0.947, p<0.0001), arm skin tissue (Spearman ρ = 0.928, p < 0.0001) and leg skin tissue (Spearman ρ = 0.870, p=0.0002) (Figure 2A). Parasite density estimates per microliter of blood or tissue were generally lower in the skin tissue compared to blood samples (Figure 2B) and not significantly different between venous or finger prick blood (p≥0.121) or between leg skin tissue or arm skin tissue (p≥=0.116). The same RNA aliquots were also processed for analysis by Nanostring expression array, a highly sensitive probe-based expression platform that we have optimized for use in *P. falciparum* (32, 33). Using a previously defined stage-specific marker set for asexual rings and mature gametocytes (33, 34), there was no evidence for higher gametocyte transcripts in skin samples compared to blood samples (Figure 2C). The two approaches to quantify gene expression also showed a strong correlation for *sbp1* and *Pfs25* (Figure 2D).

**Figure 2.**
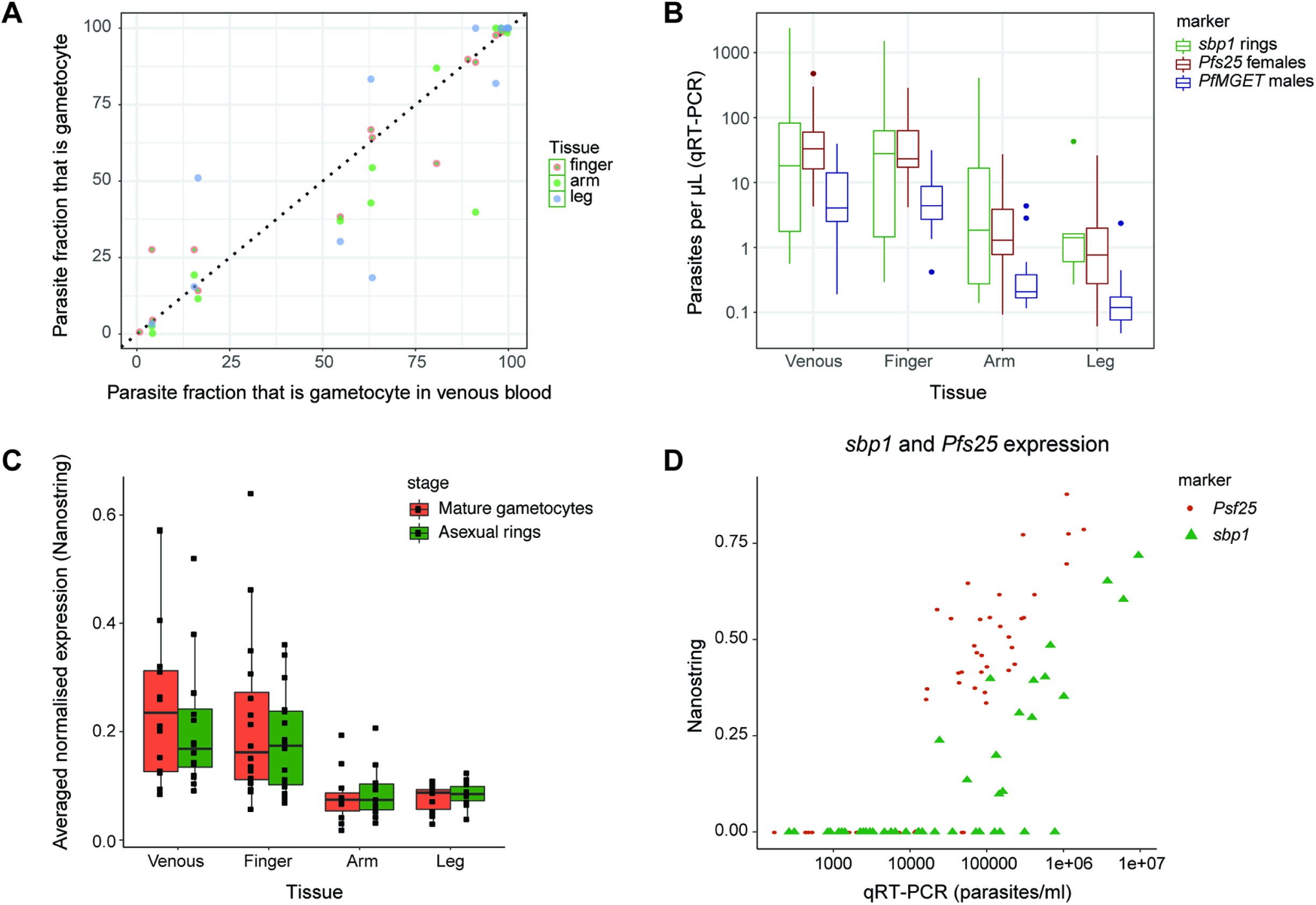
qRT-PCR and Nanostring comparison of parasite densities in skin biopsy samples and blood samples. **A.** Gametocyte fractions (the proportion of gametocytes in the total parasite biomass assessed by *sbp1*, *Pfs25* and *PfMGET* qRT-PCR) across compartments. **B-D.** Relative numbers of asexual parasites and gametocytes in skin tissue from the arm, skin tissue from the leg, finger prick and venous blood based on qRT-PCR (**B**) and Nanostring (**C**). Nanostring data were normalized on the basis of background subtraction and expression of housekeeping genes. **D.** Correlation between estimates of ring-stage asexual parasites by *sbp1* and female gametocytes by *Pfs25* for qRT-PCR (X-axis) and Nanostring (Y-axis) showing good agreement but higher sensitivity of qRT-PCR.

To directly detect gametocytes in subcutaneous tissue, skin biopsy samples that were stored in formalin were processed for imaging. Given the low densities of gametocytes predicted based on the qRT-PCR quantification (estimated median of 55.0 gametocytes in arm tissue samples (IQR 28.2-153.0) and 36.9 gametocytes in leg tissue samples (IQR 11.6-98.3); we established a protocol to image 10μm sections by confocal microscopy, hence maximizing the detectability of sparse gametocytes (Figure 3A). Skin sections were initially analysed by haematoxylin and eosin staining and labelled with the endothelial marker CD31 (Figures 3B) to confirm integrity of the tissue. Evaluation of gametocyte markers identified Pfs16 antibodies (6, 35) as highly specific and sensitive using the confocal imaging protocol (Figure 3C), while antibodies against Pfs48/45 and Pfs230 were unable to detect gametocytes in formalin fixed parasites and therefore not evaluated further. Screening of at least 12 sections per skin snip in arm and leg samples from each participant identified several putative gametocytes. A Pfs16 positive cell with a characteristic crescent shape, three-dimensional structure and nuclear stain is shown in close association with a vessel (Figure 3D and Supplementary movies 1 and 2). Based on these results, with low success gametocyte detection rates by this highly sensitive fluorescence microscopy protocol, no further gametocyte carriers were recruited as tissue donors.

**Figure 3.**
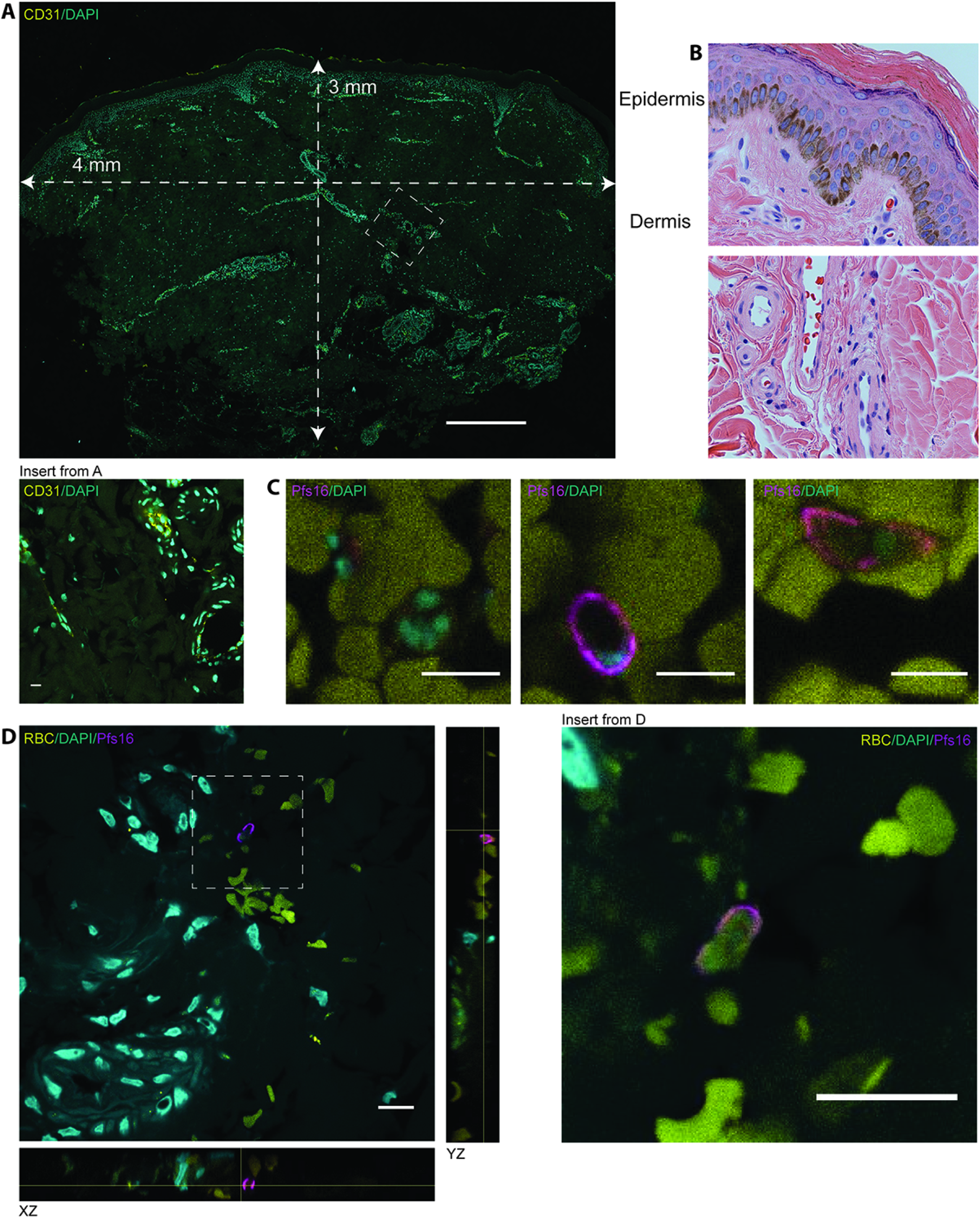
Histological analysis of skin samples. **A.** 10μm cross section of a skin snip from leg with dimensions indicated. Sample was stained with CD31 and DAPI and a maximum projection across the depth of the section is shown. The insert represents a small section including several vessels stained with CD31. Scale bar = 500μm, insert = 10μm. **B.** 3μm section of a skin snip from arm stained with haematoxylin and eosin. Sections in A and B show the different layers of the epidermis on top, followed by the dermis with multiple vessels. **C.** Samples were stained with DAPI (cyan) and Pfs16 (magenta) for gametocytes. Representative images of asexual parasite (left), an immature (middle) and mature (right) gametocyte images from control blood clots. Scale bar = 10μm. **D.** Representative image of a gametocyte in skin samples from arm. DAPI staining indicates several vessels in the vicinity of a gametocyte stained with Pfs16. XZ and YZ orientations are included to demonstrate the three-dimensional nature of the tissue section and the gametocyte. Scale bar = 10μm.

## DISCUSSION

Here, we tested a long-standing hypothesis of *P. falciparum* gametocyte sequestration in skin tissue in two populations of naturally infected individuals in Burkina Faso. By combining mosquito feeding assays and direct quantification of parasite populations in skin tissue, mosquito blood meals and blood compartments, we conclude that there is no evidence for significant skin sequestration of mature gametocytes.

Parasite sequestration in skin tissue is an intuitive explanation for how vector-borne parasites can maximize the likelihood of update by blood-feeding insects. This phenomenon, well demonstrated for a range of helminths (15–18) and protozoic trypanosomes (14), has remained speculative for *Plasmodium* parasites (22). Two recent studies in Cameroonian parasite carriers that used microscopy as diagnostic tool yielded conflicting results: one observed higher *P. falciparum* parasite prevalence in finger prick capillary blood compared to venous blood from hospital patients (36), the other found no differences for asexual parasites or gametocytes in gametocyte carriers (37). The utility of finger prick blood to estimate parasite biomass in skin tissue is uncertain. Studies published in the 1940s and 50s reported superiority of skin scarification as compared to finger prick blood samples for parasite detection (19, 20, 38). In the most extensive of these studies, in 1243 natural infections, 1 cm^2^ skin of the scapular region was very slightly scarified with 4-5 light incisions, expressing a mixture of dermal fluids and capillary blood, with the first drop appearing richest in parasites (20). This study demonstrated a 10-20% increase in prevalence of asexual parasites and gametocytes of *P. vivax, P. malariae and P. falciparum* but not *P. ovale.* Also parasite density, expressed as parasites per 15,000 examined white blood cells, appeared increased (20). In the current study, we therefore not only collected venous blood and finger prick blood but we also directly quantified parasite stage composition in skin tissue of naturally infected donors and in blood meals of mosquitoes that naturally fed on the skin of the corresponding donor. We used the absolute quantity of gametocytes and the fraction of the total parasite biomass that is gametocyte as indicators of sequestration. In skin biopsy samples, we only sporadically encountered gametocytes by histology. We chose a fluorescence imaging protocol to image thick sections by confocal microscopy. This method allowed capturing of entire parasites and three-dimensional reconstruction of parasite and surrounding tissues. Using Pfs16 labelling we classified gametocytes by crescent shape, three-dimensional structure (as opposed to non-specific speckles and autofluorescence, which is an inherent issue of this approach), nuclear stain and presence of a surrounding red blood cell. The frequency of immunofluorescence-detected gametocytes in our tissue samples was lower than that by molecular methods in a tissue sample taken during the same visit. The quality of the skin tissue, tested by analysing the tissue sections by haematoxylin and eosin staining, as well as by labelling for endothelial cells, clearly indicates they were processed and preserved well.

In contrast, molecular detection of gametocytes was successful for all tissue samples by qRT-PCR and for the majority of samples by Nanostring. Because the volume of blood is unknown in tissue samples and specifically gametocytes are hypothesized to be enriched in skin tissue (19, 20, 22), we compared the gametocyte fraction between different blood compartments and found no evidence for a biased gametocyte fraction. Gametocyte quantification in mosquito blood meals corroborated this finding and allowed a direct comparison of parasite densities. Again, we observed no evidence for higher concentrations of gametocytes in mosquitoes that fed directly on the skin of gametocyte donors compared to venous blood and observed a very strong association between gametocyte fractions from the different blood compartments. There must therefore be an alternative explanation for the higher infection rates that we, in line with other studies (9, 10), observed in direct skin feeding experiments compared to membrane feeding experiments using venous blood. Gametocyte activation may occur following phlebotomy and may reduce infection rates observed following membrane feeding. In addition, anticoagulants used in phlebotomy can have a pronounced effect on mosquito infection rates (39). Although heparin is the preferred anticoagulant (39), it may still have a disadvantageous impact on sporogonic development. In malaria-naïve individuals in whom *P. falciparum* gametocytes were induced during controlled human malaria infection studies, replacement of heparin plasma by serum resulted in increased mosquito infection rates (10). Since human immune responses are unlikely to be of relevance in these gametocytaemic volunteers, this observation provides additional indirect evidence for a transmission modulatory effect of heparin.

We conclude that there is no evidence for gametocyte sequestration in skin tissue. Our findings argue against a long-standing hypothesis that never had a solid evidence base or proposed mechanism. Since the deformability of erythrocytes infected with mature gametocytes is similar to that of uninfected erythrocytes (23, 40) and there is no evidence for antigens on the surface of mature gametocyte-infected erythrocytes (41, 42), it is perhaps unsurprising that gametocyte concentrations are similar in the different blood compartments. While direct skin-feeding assays tend to result in higher infectivity compared that observed in indirect feeding procedures using venous blood, our data demonstrate that any differences observed are based on technical rather than biological differences in the feeding procedure. Our findings also indicate that gametocyte levels in venous or finger prick blood can be used to predict onward transmission potential to mosquitoes. Our findings thus pave the way for methodologies to quantify the human infectious reservoir based on conventional blood sampling approaches to support the deployment and monitoring of malaria elimination efforts for maximum public health impact.

## MATERIALS AND METHODS

### Ethics statement

Ethical approval for the studies was granted by the Ethical Review Committee of the Ministry of Health of Burkina Faso (Deliberation numbers 2016-03-033 and 2017-02-018) and the Ethics Committee of the London School of Hygiene and Tropical Medicine (#10489 and #11962). Individual written informed consent was obtained from each participant prior to enrolment. Malaria cases were treated according to the National guidelines in Burkina Faso (43).

### Study site and population

Study participants were recruited in the village of Balonghin, located in Saponé district, in Burkina Faso. Malaria transmission is seasonal and intense. The main malaria vectors are *Anophele gambiae s.s*, *An. coluzzii, An. arabiensis* and *An. funestus*. *P. falciparum* parasite carriage and gametocyte carriage by molecular methods in the study area are 51-84% and 49-75%, respectively (44).

### Study design

#### Paired skin feeding and membrane feeding study

This study was conducted in October-December 2017. Individuals from the eligible age range (15-50 years) in the study area were invited to study information meetings based on a village census list and, if expressing an interest to participate, invited for screening at Balonghin health facility. Eligible participants had *P. falciparum* gametocyte densities ≥1 gametocyte/500 leucocytes by microscopy (≥16 gametocytes/μL when assuming 8000 leucocytes/ μL). Exclusion criteria were: signs of acute or chronic disease that required immediate clinical care; haemoglobin concentration <8 g/dL; current or previous participation in malaria vaccine trials; recent blood transfusion or administration of blood products; use of antimalarials in the last 2 weeks; co-infection with *P*. malariae or *P*. *ovale*. Eligible participants were provided transport to the Centre National de Recherche et de Formation sur le Paludisme (CNRFP) in Ouagadougou for membrane feeding and skin feeding. Immediately after venipuncture in lithium heparin and EDTA tubes (BD Vacutainer™), 400-500µL of heparinized blood in duplicate (for infectivity) and 400-500µL EDTA blood (for gametocyte quantification in blood meals) was offered to 60 starved 4–5-day-old female *An. coluzzii* mosquitoes via an artificial membrane attached to a water-jacketed glass feeder maintained at 37°C (28). After exactly 15 minutes of feeding in the dark, fully fed mosquitoes from heparin blood were transferred to storage cups by aspiration and maintained with glucose solution at 27-29°C for 6-8 days before dissection with 1% mercurochrome staining and examination for oocysts by two independent microscopists. From mosquitoes that fed on EDTA blood, 16 fully fed mosquitoes were sacrificed after feeding for exactly 15 minutes by sharp needle puncture of their midguts to release the blood contents into 50µl of RNAprotect cell reagent; blood meal material was stored for individual mosquitoes at −80°C. Immediately following membrane feeding, direct skin feeding took place. The participant’s calves were exposed to 60 mosquitoes distributed over 2 paper cups that were allowed to feed for exactly 15 minutes. From this group, 12 fully fed mosquitoes were immediately sacrificed and their midguts punctured as described above. Remaining mosquitoes were maintained on glucose solution before dissections for oocyst presence, as above. In addition to the membrane and direct skin feeding assays, K2EDTA blood was collected by venipuncture (BD Vacutainer™) and finger prick (BD Microtainer®).

### Skin biopsy study

In the period September 2016-March 2017, adults (aged 18-50 years) were invited for study participation as described above. Participants were eligible if they had *P. falciparum* gametocyte densities ≥1 gametocyte/500 leucocytes by microscopy (≥16 gametocytes/μL). For skin biopsy sampling, exclusion criteria were signs of acute or chronic disease that requires immediate clinical care; haemoglobin concentration <11 g/dL; skin infections or conditions; history of vasovagal responses to blood sampling or biopsies; allergy to lidocaine/ prilocaine. Eligible individuals were invited to the CNRFP central lab in Ouagadougou on two occasions, 4 days apart for sample collection. At each occasion, skin biopsy samples including the dermis and hypodermis were taken from under the arm (n=2) and leg (n=2) using single use punchers (4mm Biopsy Punch; Miltex Inc. York, US). This procedure was performed 1 hour after applying local anaesthetic by means of a xylocaine-adrenaline by a qualified dermatologist. Half of the biopsy samples (one each from arm and leg) were immediately immersed in 2 mL of 10% formalin and placed at 4°C overnight; following washing, samples were stored in 2 mL of 70% ethanol and stored at 4°C until further processing. Other biopsy samples were transferred to 1000 µL RNALater stabilization reagent (Qiagen), incubated overnight at 2-8°C and then transferred to −80°C. Finger prick and venous blood samples were collected in EDTA-coated tubes, as above.

### Molecular analysis

Mosquito homogenates were pooled (4 mosquitoes in a total of 200µl of RNAprotect per pool) with 4 pools (16 mosquitoes) for membrane feeding experiments and 3 pools per skin feeding experiment (12 mosquitoes). Mosquitoes where no blood was released into RNAprotect (upon visual expectation upon thawing) were not used for extraction and, as a result, fewer pools of mosquitoes were extracted. Nucleic acids from these 200μL mosquito pools and from 100μL venous and finger prick whole blood samples in RNAprotect Cell Reagent were isolated using the bead-based MagNAPure LC automatic extractor (Total Nucleic Acid Isolation Kit—High Performance, Roche Applied Science) and eluted in 50μL of water. In these samples, ring-stage asexual parasites, female gametocytes and male gametocytes were quantified by individual quantitative reverse-transcription PCR (qRT-PCR) assays targeting *sbp1* (31); *Pfs25* (45) and *PfMGET* (29), respectively. Skin biopsy samples were immediately stored in RNAlater solution after collection. RNA extraction from skin tissue was performed using the Qiagen RNeasy Plus Mini kit (Qiagen). First, the tissue samples were removed from RNAlater solution and then homogenized in RLT lysis buffer (Qiagen) using Polytron Homogenizer (Kinematica). The homogenized lysate was passed through genomic DNA eliminator columns (Qiagen) and subsequently applied to RNeasy spin columns. Following several washes, RNA was eluted in nuclease-free water according to the manufacturer’s instructions.

The NanoString nCounter custom code set included differentially expressed genes to distinguish specific *P. falciparum* parasite stages as defined from our previous study (34). A total of 456 parasite genes were included in the custom probe set including housekeeping genes. 161 genes representing asexual circulating stages, 147 genes representing asexual sequestering stages, 26 genes representing gametocyte rings, 27 immature gametocytes and 29 mature gametocyte genes. The remaining set was not annotated for any of these parasite stages. For NanoString analysis, 5 μl of purified total RNA was used for initial hybridization reaction. RNA from each sample was allowed to hybridize with reporter and capture probes at 65°C for 20 hours according to the nCounter gene expression assay protocol (NanoString Technologies). RNA-probe complexes were immobilized to nCounter cartridge followed by scanning in the nCounter Digital Analyzer. Data was first normalized by applying background subtraction and then normalized to expression of housekeeping genes using the R package “NanoStringNorm”. The dataset was then quantile normalized using the R package “aroma.light” and rank scaled. Mature gametocyte and asexual marker genes, as defined in^33^, were then averaged per patient, per tissue and per visit.

### Histological analysis of skin samples

Skin biopsies were processed by passing through an increasing alcohol gradient and xylene before embedded in paraffin wax. 10μm sections of biopsy samples were cut on a microtome and placed on adhesion slides (SuperFrost® Plus Gold, VWR). Slides were dried at room temperature for at least one hour then baked overnight at 42°C. The slides were allowed to reach room temperature before proceeding with the staining protocol. Slides were incubated at 60°C to melt the wax around the section; sections were cleared with xylene and rehydrated by passing through a decreasing alcohol gradient (xylene: 5 minutes twice; 100% ethanol: 3 minutes, twice; 90% ethanol: 3 minutes, twice; 70% ethanol: 3 minutes, twice). After incubation in distilled water for 3 minutes, heat induced antigen retrieval was performed using citrate buffer pH 6.0 (TCS Biosciences) in a table top autoclave. Slides were immersed in buffer using a metal rack in an empty tip box (without lid) and autoclave initiated until it reached 126°C, at which point the autoclave was unplugged and slides allowed to incubate in the autoclave for a further 10 minutes. Subsequently, the slides were removed and cooled in their buffer in a running water bath. Once at room temperature, slides were transferred to distilled water and then TBST (Tris Buffered Saline with 0.05% Tween 20) for 3 minutes each. Slides were then blocked with goat block containing 2.5% normal goat serum (Vector Laboratories) complemented with 2.5% normal human serum (ThermoFisher Scientific). All blocking and staining were performed in a humidified chamber. All staining solutions were removed by tapping the side of the slide gently on tissue paper. Excess liquid was removed by wicking away with tissue paper, being careful not to touch the sections. This was done to maintain intact, well-formed skin sections which are particularly delicate. After 30-60 minutes blocking at room temperature, the slides were incubated in primary antibodies diluted in goat block. Sections were stained with 1:20 (1.12μg/ml) mouse anti-CD31 (Cell Marque: clone JC70) at 4°C overnight or 1:1250 (1.04μg/ml) rabbit anti-Pfs16 (6) at room temperature for one hour. The slides were then washed with TBST for 3 minutes thrice before adding 1:100 goat anti-mouse IgG-AlexaFluor488 (ThermoFisher, A-11029) or 1:250 goat anti-rabbit IgG-AlexaFluor647 (ThermoFisher, A-21245) secondary antibody diluted in goat block and incubated at room temperature for 30 minutes. Following secondary antibody staining, the sections were washed twice with TBST and then once with TBS for 3 minutes each, before incubation with 2.5nM final concentration of DAPI diluted in TBS for 10 mins at room temperature. Sections were washed twice more in TBS for 3 minutes, before addition of TrueView autofluorescence quenching reagent (Vector Laboratories) and incubation for 3 minutes at room temperature. Sections were washed once more in TBS for 5 minutes before mounting with Vectashield Vibrance mountant (Vector Laboratories). Slides were viewed on a Nikon A1R inverted confocal microscope with Piezo Z-drive to acquire z-stacks. In addition to skin biopsies, clots of cultured *P. falciparum* parasites (strains Pf2004, 3D7 and NF54) were generated to act as positive and negative controls. Asexual and mixed asexual-immature gametocyte clots and mature gametocyte clots were generated as described previously (6). Sections of formalin fixed paraffin embedded blocks were used to optimise Pfs16 antibody and DAPI staining and determine the staining of mature gametocytes. Using these controls gametocytes in the skin were determined by their circumferential staining with Pfs16 and obvious outline of a red blood cell. Red blood cells were determined by their bright autofluorescence under 488nm laser light. Images and movies were generated using Image J software.

### Sample size justification

For the paired skin feeding-membrane feeding study, we assumed an average of 15% infected mosquitoes in patent gametocyte carriers with a standard deviation of 20% and a within subject correlation of the outcome of 0.5 (9, 46, 47). If we then expected two-fold higher mosquito infection rates in direct skin feeding, 17 paired membrane feeding and skin-feeding experiments on patent gametocyte carriers would give 80% power to detect this difference at an alpha of 0.05. Sample size justification for skin-biopsy sampling was based on a paired comparison of the proportion of the total parasite population that is mature gametocyte. We expected that 73% of the skin snip biopsy samples had higher gametocyte concentrations, based on a meta-analysis that demonstrated enhanced infectivity following skin feeding compared to venous blood membrane-feeding (9). When assuming that 70% of infected adults have detectable malaria parasites in skin tissue and allow quantification of the proportion of parasites that is gametocyte, and a lower limit of the 95%-CI >50%, 45 paired skin snip samples and venous/finger prick blood samples would give 83% power with an alpha of 0.05 to detect a different in parasite stage composition. A go/no-go criterion was defined where an initial 10 gametocyte carriers were recruited for biopsy samples and additional participants would only be recruited if gametocytes were detected in ≥50% of all samples.

### Statistical analysis

All statistical analyses were performed in STATA version 15.0 (Statacorp; College Station, TX, US). The proportion of infectious gametocyte carriers was compared between paired feeding experiments using McNemar’s test; the proportion of infected mosquitoes was compared between direct skin feeding and membrane feeding using logistic regression controlling for study participant as a fixed effect. Spearman non-parametric correlation coefficients were calculated to assess associations between continuous variables; the paired Wilcoxon rank-sum test was used to compare parasite densities between blood or tissue samples from the same participants. The gametocyte fraction was calculated as the sum of male and female gametocytes, expressed as a proportion of the total parasite biomass of asexual ring-stage parasites and gametocytes.

## Supplemental data

### Supplementary movie 1 (3D movie)

3D projection of Z-stack of mature gametocyte in skin snip. This movie shows the 3D reconstruction of the z-stack (step-size 0.2 micron) to illustrate the localisation of a mature gametocyte. The gametocyte is stained with Pfs16 (magenta), denoted by DAPI staining (cyan), and within an RBC (yellow. It is in close proximity to skin vasculature. Movie was generated using Image J software.

### Supplementary Movie 2 (Z stack)

Z-stack of mature gametocyte in skin snip. Confocal z-stack of mature gametocyte taken across the whole thickness of the section (step-size 0.2 micron). Gametocyte stained with Pfs16 (magenta), with DAPI (cyan) nuclear staining. Movie generated using Image J software.

## Data availability

Data underlying this manuscript are available through https://datadryad.org/stash/share/_Di1z3S3jl2ahewKXHAXHfAtl7slSBGNAZmgueslqbI.

## Acknowledgements

We would like to thank all study participants from Balonghin, Burkina Faso, for their participation. We further thank Fiona McMonagle for her guidance and assistance in the histology work.

